# Arabidopsis ecotype screening reveals novel sources of clubroot resistance and insights into resistance inheritance

**DOI:** 10.1101/2025.04.15.649024

**Authors:** Melaine Gonzalez-Garcia, Soham Mukhopadhyay, Ian Major, Edel Pérez-Lopez

## Abstract

Clubroot, caused by *Plasmodiophora brassicae*, poses a persistent threat to Brassicaceae crops, particularly in regions where resistant cultivars are under strong selection pressure. To identify new sources of resistance and better understand the underlying genetic mechanisms, we evaluated 60 *Arabidopsis thaliana* ecotypes against the highly virulent Canadian pathotype 3A. Using stringent phenotyping criteria, pathogen DNA quantification, and survival analysis, we identified eight resistant ecotypes, including two novel sources, Marce-1 and DraII-6. DraII-6 exhibited exceptionally low disease symptoms and a high survival rate. While the resistance gene RPB1/WeiTsing was present in most ecotypes, its expression in DraII-6 was significantly elevated at early infection stages, suggesting a potential role in pathogen suppression. However, genetic analysis of F1 and F2 progeny from a DraII-6 × Col-0 cross revealed a recessive resistance pattern, supporting the hypothesis that RPB1 alone may not be sufficient to confer resistance to clubroot in DraII-6. Our findings highlight the complexity of clubroot resistance and the need for further research into gene regulation and resistance networks beyond RPB1, particularly in the context of translating Arabidopsis-based insights to Brassica crops.

**GRAPHICAL ABSTRACT:** 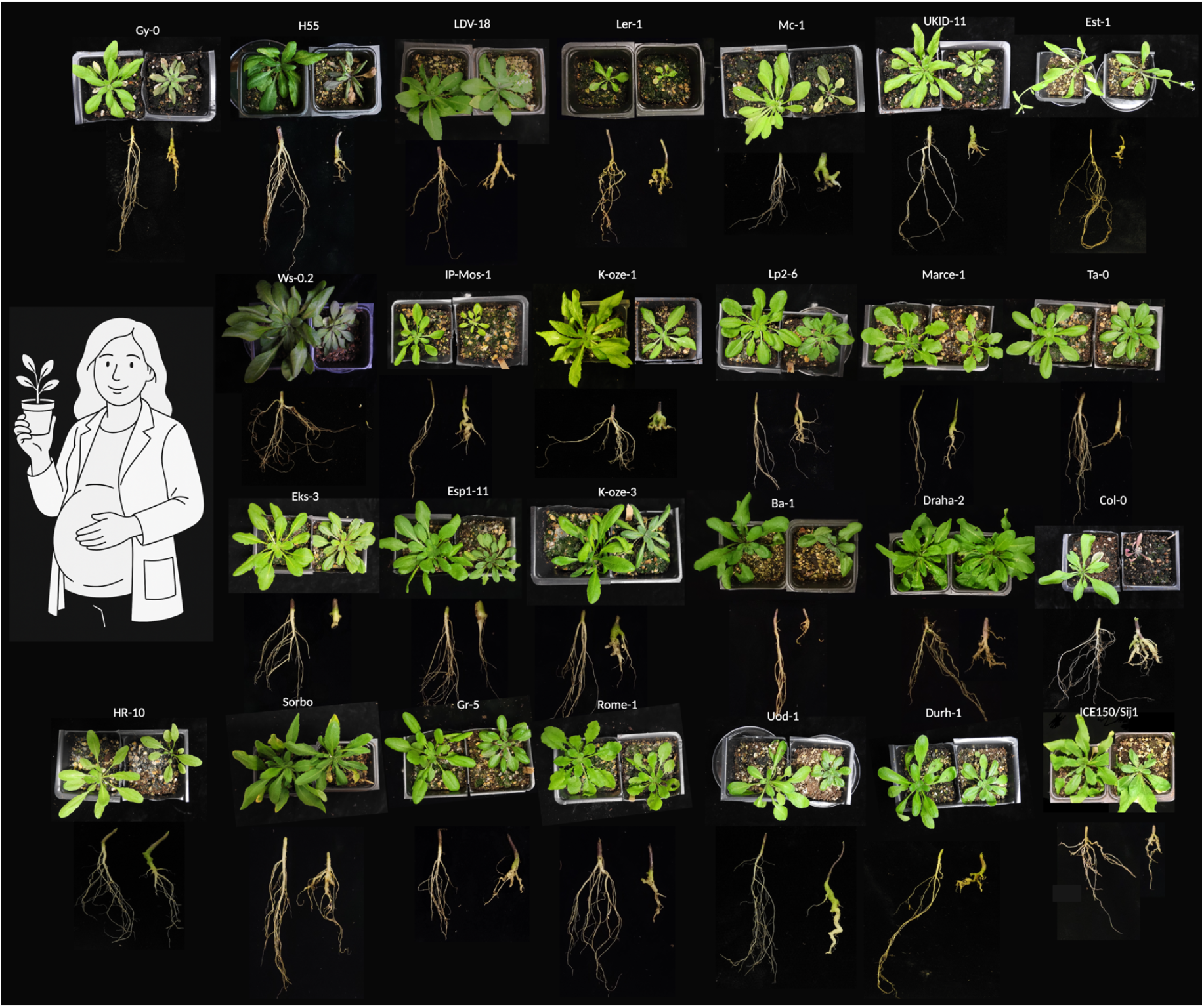

Natural variability of *Arabidopsis thaliana* ecotypes in response to the Canadian clubroot pathotype 3A. The abstract also celebrates the first author, Melaine Gonzalez-Garcia, who is submitting this manuscript just one week before welcoming her first child.

---

Clubroot, caused by the obligate biotrophic pathogen *Plasmodiophora brassicae*, is a devastating disease of cruciferous crops, leading to global yield losses of 10–15% (Javed et al., 2022). This soil-borne pathogen induces gall formation on infected roots, disrupting water and nutrient uptake, and ultimately causing stunting, wilting, and plant death (Dixon, 2009; Botero et al., 2019). *P. brassicae* has a complex life cycle, including long-lived resting spores that can remain viable in the soil for up to two decades (Dixon, 2009). Disease symptoms result primarily from secondary infection, during which the pathogen hijacks host developmental pathways to induce abnormal cell proliferation and vascular distortion (Malinowski et al., 2012; 2019).

Currently, clubroot management remains a significant challenge. While preventive strategies such as crop rotation, soil amendments, and chemical treatments can reduce disease severity (Peng et al., 2014), the deployment of clubroot-resistant (CR) cultivars remains the most effective and sustainable control method (Cao et al., 2014; Mukhopadhyay et al., 2024). Since the introduction of the first CR canola cultivar in Canada in 2009 (Deora et al., 2015), the reliance on genetic resistance has grown. However, this strategy has accelerated the emergence of virulent pathotypes capable of overcoming resistance, particularly under the selection pressure of widespread CR cultivar use (Strelkov et al., 2018; Sedaghatkish et al., 2019).

This evolutionary arms race underscores the need for continuous discovery of new resistance sources. Numerous resistance (R) loci have been identified in *Brassica rapa* (e.g., *Crr1, CRa, CRb, CRk*) and *B. oleracea* (Alamery et al., 2018; Yang et al., 2022), but *Arabidopsis thaliana* has also proven to be a valuable model for dissecting host-pathogen interactions and identifying resistance genes (Alix et al., 2007; Jubault et al., 2008). Several surveys using *A. thaliana* ecotypes have revealed natural variation in resistance against various *P. brassicae* pathotypes (Fuchs and Sacristan, 1996; Alix et al., 2007; Jubault et al., 2008; Ochoa et al., 2023; Wang et al., 2023).

A recent study characterized *RPB1*, also known as WeiTsing, a broad-spectrum resistance gene on chromosome 1 of *A. thaliana*, originally identified in the Est-1 background (Ochoa et al., 2023; Wang et al., 2023). Despite the excitement around this gene, their validation was based solely on infection assays using CR ecotypes against six Chinese *P. brassicae* isolates and a European clubroot pathotype. This narrow scope raises concerns about the true breadth of resistance conferred by *RPB1/WeiTsing*, particularly against diverse global pathotypes, including those prevalent in Canada, a major canola producer. Here, we evaluated the resistance of 60 *A. thaliana* ecotypes against the 3A pathotype of *P. brassicae*, one of the most widespread and virulent pathotype in Canada (Storfie et al., 2025).

The *A. thaliana* ecotypes screened in this study included several previously identified as CR in Europe (EUR; Ochoa et al., 2023) and China (CHN; Wang et al., 2023): Est-1, Tsu-0, Durh-1, HR-10, Uod-1, and Pub2-23. Additionally, we included susceptible ecotypes such as Ta-0, LP2-6, Sorbo, Ler-0, Ler-1, Duk, and Col-0 (Fig. 1, Table S1). The remaining ecotypes were primarily selected based on their geographic origin, with an emphasis on Eastern Europe, considered the center of origin for clubroot (Fig. S1) (Javed et al., 2022).

**Figure 1.**
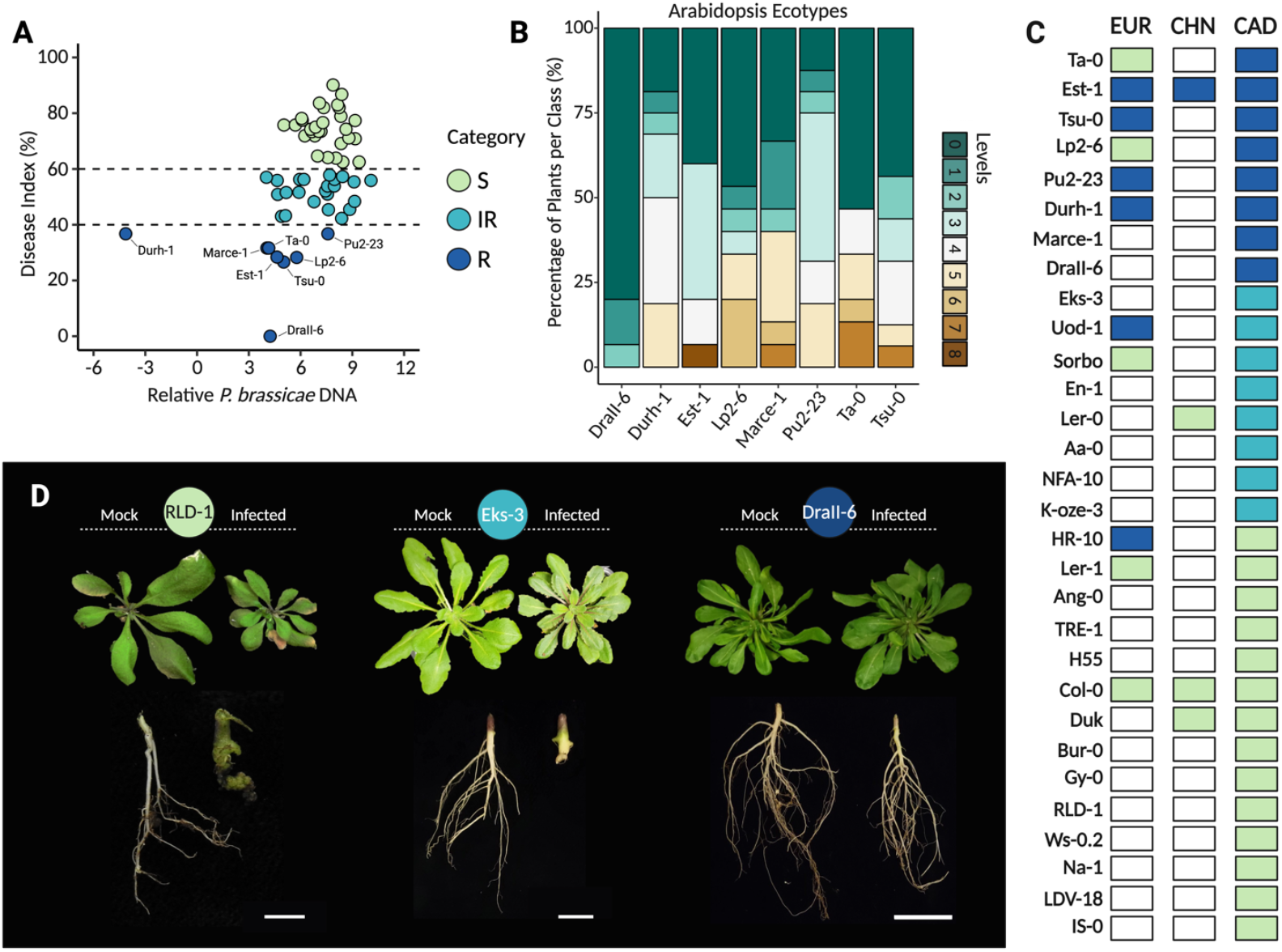
Natural variation in disease index and pathogen load across *Arabidopsis thaliana* ecotypes in response to *Plasmodiophora brassicae* pathotype 3A infection. A) Relationship between disease index and relative P. brassicae DNA content in 60 ecotypes categorized as resistant (R = 0% to 40% DI in green), intermediate-resistant (IR = 41% to 60% DI in light blue), or susceptible (S = 61% to 100% DI in dark blue). A Spearman correlation was performed to assess the association between relative *P. brassicae* DNA levels and disease index (r = 0.33; P = 0.009). Values plotted represent the mean of three biological replicates for DI and three technical replicates for P. brassicae DNA titer. Statistical analyses were conducted in R (version 4.4.2) and RStudio (version 2024.12.1+563) using the packages stats, survival, and graphics, with visualizations generated using ggplot2. B) Percentage of plants per disease severity class (0–8) among CR ecotypes. Table S2 provides a comprehensive overview of the disease scoring criteria and corresponding severity levels. C) Comparison of Arabidopsis ecotype responses to the clubroot pathogen across studies from Europe (EUR), China (CHN), and the present study using a Canadian (CAD) isolate. Only 30 ecotypes are shown, including the 13 that were evaluated in both this study and at least one previous study. The rest are presented in Table S1 and S4. D) Representative root and shoot phenotypes of RLD-1 (S), Eks-3 (IR), and DraII-6 (R) at 21 days post-inoculation. Scale bar = 1 cm.

While the standard disease rating system uses a 0–4 scale, where 0 indicates no symptoms and 4 indicates severe galling, root deformation, and impaired growth, we employed a more comprehensive, cumulative scale (Table S2). This expanded scale allowed for detailed characterization of both root and shoot symptoms, distinguishing eight levels of infection. These were then used to calculate the disease index (DI) using the established formula (Salih et al., 2024). Relative *P. brassicae* DNA levels were also calculated for each ecotype (Table S3) and plotted alongside the DI to classify ecotypes as resistant, intermediate-resistant, or susceptible.

Consistent with the EUR and CHN screenings, our study conducted against a Canadian pathotype (CAD) confirmed that Est-1 exhibits broad-spectrum resistance against multiple *P. brassicae* isolates, including P1+ from Europe, six field isolates from China, and the 3A pathotype (Fig. 1). However, no other ecotype was consistently classified as resistant across all three studies. While Tsu-0, Pu2-23, and Durh-1 each displayed a DI below 40%, the threshold we used to define resistance (Fig. 1A–C), the *P. brassicae* DNA titer in Tsu-0 and Pu2-23 was comparable to the titer of some ecotypes classified as intermediate-resistant or susceptible (Fig. 1A). This disconnection between disease symptoms and pathogen titer has been previously reported (Ochoa et al., 2023), although its exact cause remains unclear. One possible explanation is the clubroot pathogen’s ability to manipulate host hormone signaling, particularly cytokinins and auxins, which may decouple symptom severity from actual pathogen load (Javed et al., 2022; Vañó et al., 2023). This supports the idea that clubroot pathogenesis is highly complex, where even a relatively low pathogen burden can induce severe symptoms through interference with host hormonal homeostasis (Vañó et al., 2023). Another notable inconsistency among the studies is the identification of Ta-0 and Lp2-6 as clubroot-resistant (CR) against the Canadian pathotype 3A, despite both being susceptible to the EUR Pb1+ isolate (Fig. 1A, C). Conversely, Sorbo and Ler-0, previously classified as susceptible in the EUR and CHN studies, were identified by us as intermediate-resistant, with a DI between 40% and 60% (Fig. 1A, C). The differences observed among studies can be attributed to variations in the pathogenicity of the pathotypes and isolates tested by each group, as well as differences in inoculum concentrations and the thresholds used to define CR (Ochoa et al., 2023; Wang et al., 2023). However, considering that our parameters were the most stringent, using a lower DI threshold and a higher concentration of *P. brassicae* spores, we are confident in the robustness of our results.

In total, eight *A. thaliana* ecotypes were identified as CR in our study, including two reported here for the first time: Marce-1 and DraII-6 (Fig. 1A–D, Table S4). In terms of pathogen DNA titer, both ecotypes exhibited similar levels (Fig. 1A, Table S4). However, DraII-6 stood out among all CR ecotypes, with a DI close to zero (Fig. 1A, Table S4). This was consistently evident during phenotyping, as DraII-6 plants showed virtually no visible symptoms, neither in the shoots nor in the roots, except for slight thickening of the main root (Fig. 1D). These subtle signs were only noticeable when compared to susceptible or intermediate-resistant ecotypes at 21 dpi (Fig. 1D). Given that resistance in at least six of the eight CR ecotypes has been attributed to the RPB1 gene, we screened all 60 ecotypes for its presence using PCR, following the protocol described by Wang et al. (2023) (Table S3). We found that most ecotypes, including Marce-1, DraII-6, and several susceptible ones, harbored the RPB1 gene (Fig. S2, Table S5). These results are consistent with previous studies, which suggest that resistance to *P. brassicae* is not solely determined by the presence of RPB1, but rather by the regulation of its expression (Ochoa et al., 2023; Wang et al., 2023). Another element to consider that may explain the differences observed among the three studies is the variation in effector repertoires among isolates. These differences were recently highlighted by our group after completing the first telomere-to-telomere *P. brassicae* genome and investigating the evolution of the *Plasmodiophorids* secretome (Javed et al., 2024; Mukhopadhyay et al., 2025). To date, how *P. brassicae* triggers RPB1-mediated CR remains unknown, although it is expected to be effector-mediated (Ochoa et al., 2023; Wang et al., 2023).

A common limitation across all studies using Arabidopsis as a source of CR is that ecotype evaluation is typically conducted only at 21 dpi or sooner, which usually coincides with the flowering stage (Ochoa et al., 2023; Wang et al., 2023; Salih et al., 2024). This approach does not allow assessment of whether the low DI observed at that point persists through later stages of development, nor whether the plant can complete its life cycle and successfully reproduce. These traits are crucial for translating findings from Arabidopsis to economically important Brassica crops such as *Brassica napus* and *Brassica rapa*, where full-cycle productivity is key for the industry.

To determine whether the eight ecotypes identified as CR can complete their life cycle under infection, we monitored their survival up to 42 dpi, along with the susceptible control ecotype Col-0. Symptoms were recorded weekly, and plants were photographed (Fig. 2A). We conducted a survival analysis using Kaplan–Meier estimators to assess survival across the selected resistant ecotypes. Each plant was scored weekly as either 0 (alive) or 1 (dead), and survival curves were compared using the Log-Rank test with Bonferroni correction. The results showed that DraII-6, Est-1, Tsu-0, Durh-1, and Marce-1 all had significantly higher survival probabilities than Col-0 (Fig. 2B, Table S6). An important observation is that most of the resistant ecotypes are late-flowering types, which may influence the delayed onset of clubroot symptoms such as gall size, gall formation on lateral roots, and wilting. A potential link between life span and immunity in Arabidopsis has been previously proposed, showing that reduced life span is associated with decreased expression of immunity-related genes (Glander et al., 2018). This is the first time such a phenomenon has been reported in the context of clubroot disease.

**Figure 2.**
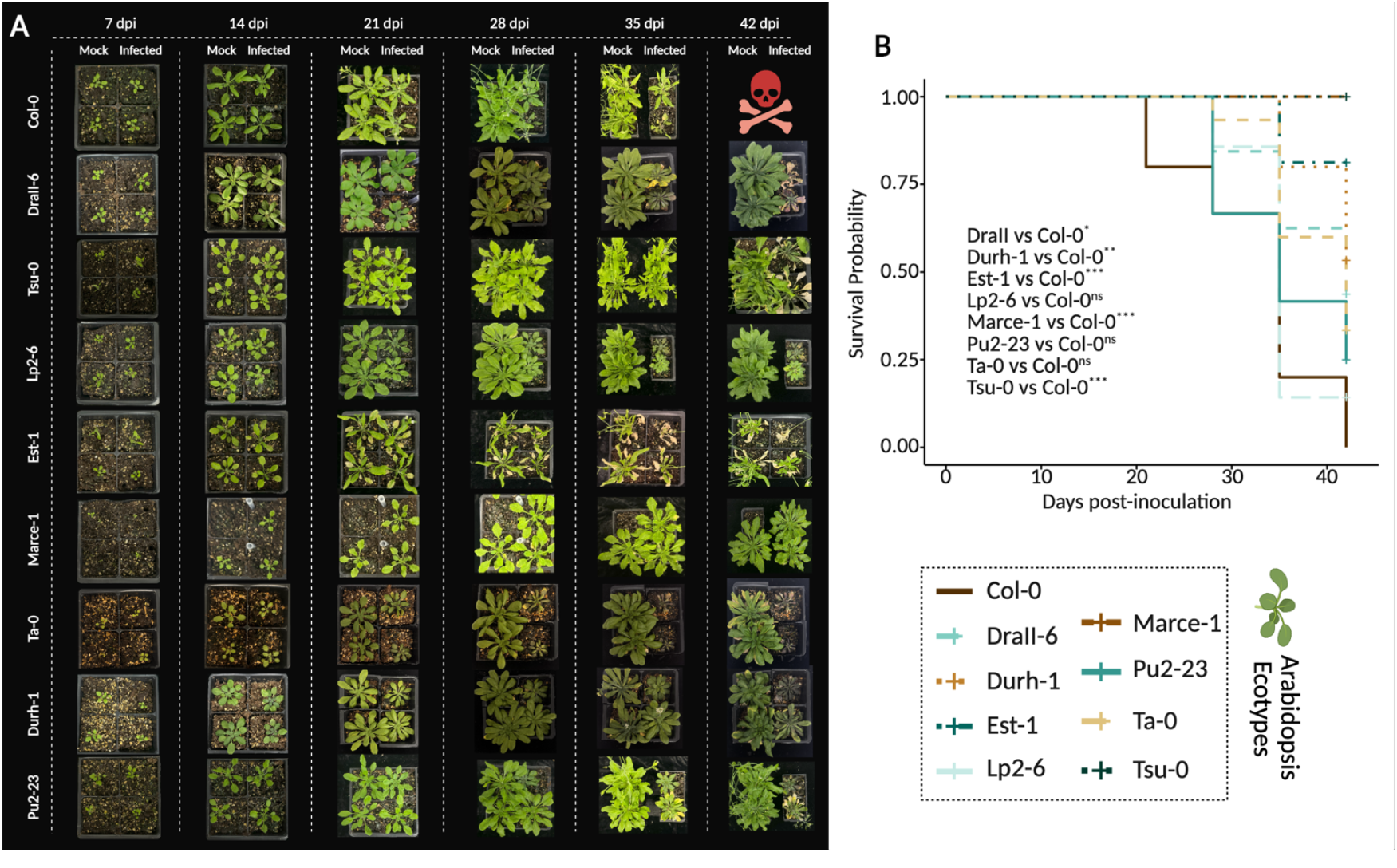
Survival analysis of CR ecotypes compared to the susceptible Col-0. A) Overview of Col-0 and eight CR ecotypes from 7 to 42 days post-inoculation (dpi), under both infected and mock conditions. Col-0 plants did not survive to 42 dpi under either condition, whereas most CR ecotypes—identified as late-flowering—remained viable. To accommodate a global view of the time course of all ecotypes, plant images were resized individually as necessary. Differences in plant sizes may not necessarily reflect actual differences. B) Survival analysis using Kaplan–Meier estimators. Each plant was scored as 0 (alive) or 1 (dead) for each evaluation day. Survival curves for each ecotype were compared using the log-rank test with Bonferroni correction (Chi-square = 73.99; df = 8; P < 0.0001). Pairwise comparisons from the log-rank test are shown in the figure. Statistical analyses were conducted in R (version 4.4.2) and RStudio (version 2024.12.1+563) using the packages stats, survival, and graphics, with visualizations generated using ggplot2.

A key question remained: Is RPB1/WeiTsing responsible for the high level of clubroot resistance observed in DraII-6? To investigate this, we analyzed the expression of RPB1 in DraII-6, comparing it to the well-characterized resistant ecotype Est-1 and the susceptible ecotype RLD-1. This was evaluated at 4- and 7-dpi, and interestingly, RPB1 expression in DraII-6 was significantly higher than in both Est-1 and RLD-1 at 4 dpi. By 7 dpi, expression levels in DraIi-6 were similar to those in Est-1 and both remained significantly higher than in the susceptible RLD-1 (Fig. S3, Table S7). These results support the hypothesis that RPB1 may play a key role in mediating the high level of clubroot resistance observed in DraII-6, particularly since similar expression levels were previously reported in Est-1 and Uod-1 during infection at 7 dpi (Ochoa et al., 2023).

Moreover, the elevated expression of RPB1 as early as 4 dpi could be especially relevant, although this early time point has not been thoroughly explored in previous studies. Early activation of RPB1 might interfere with the pathogen’s ability to establish itself in host tissue, potentially affecting the critical transition from primary plasmodia to secondary zoosporangia (Javed et al., 2022). Given that RPB1 encodes a calcium channel (Wang et al., 2023), its early expression could alter intracellular calcium homeostasis, possibly creating a hostile or toxic environment for *P. brassicae* development and triggering further immune responses. This novel observation opens the door to further investigations into the timing and mechanistic role of RPB1-mediated resistance during the initial stages of infection.

To further investigate the involvement of RPB1 in DraII-6 resistance, we studied its segregation in the F1 generation resulting from a cross between DraII-6 (female) and the susceptible ecotype Col-0 (male). The F1 progeny were phenotyped as described above. Surprisingly, despite previous reports describing RPB1 as a dominant gene (Wang et al., 2023), the F1 plants were fully susceptible to clubroot. However, the severity of the infection was significantly lower than in the Col-0 parent (Fig. 3, Table S8). To gain further insight, we phenotyped 240 F2 individuals derived from open pollination of the F1. The results showed a segregation pattern of approximately 25% resistant and 75% susceptible plants, consistent with a recessive inheritance model (Fig. 3B, Table S8). This contradicts previous findings that classified RPB1-mediated resistance as dominant, although we did not corroborate RPB1 segregation in the F2 (Fig. 3B). These findings support an alternative hypothesis: RPB1 alone may not be sufficient to confer resistance to clubroot in DraII-6. Additional genetic or regulatory factors may be involved, and further investigation is warranted to fully understand the resistance mechanism in this ecotype.

**Figure 3.**
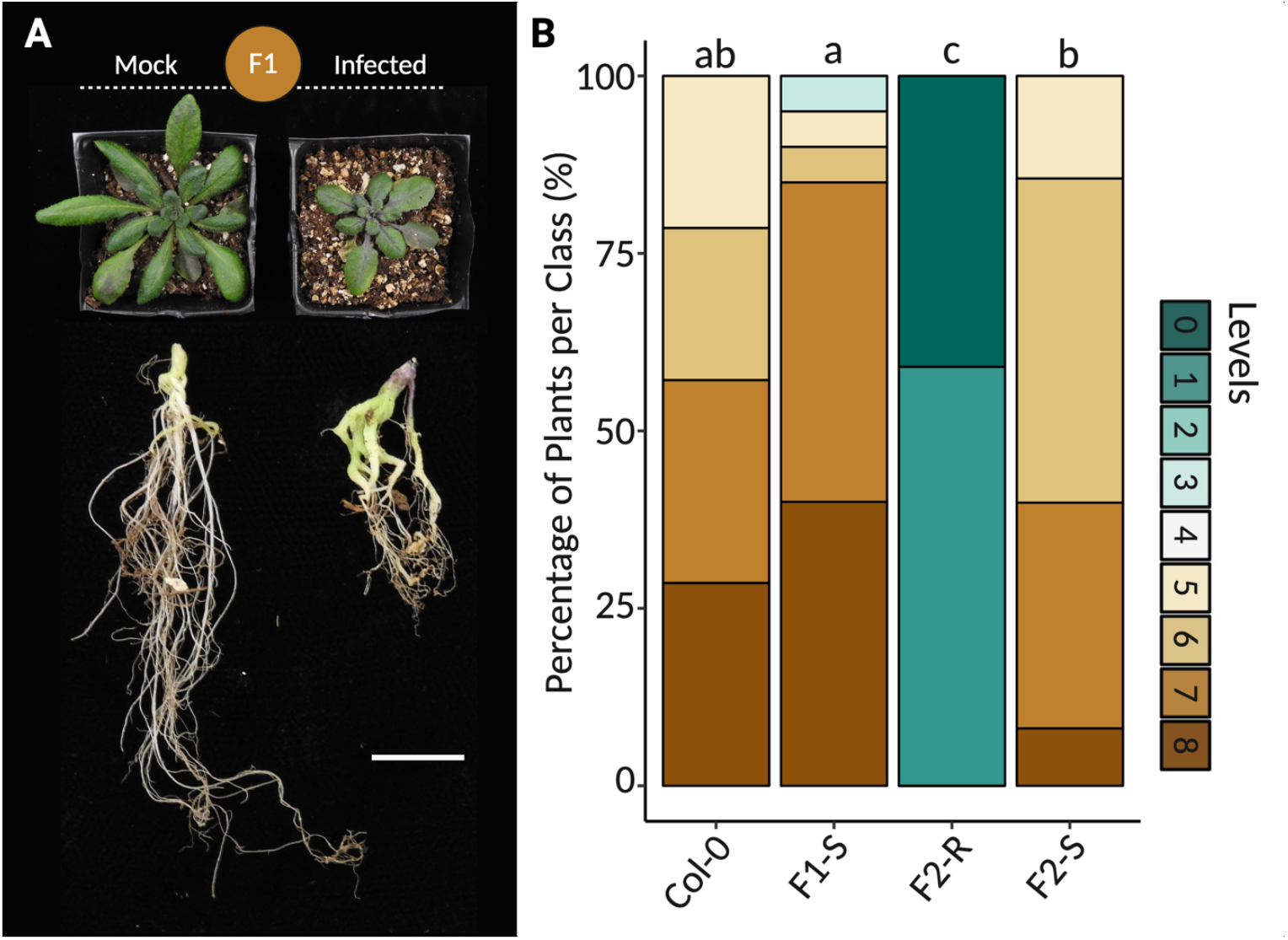
Phenotyping of DraII-6 × Col-0 cross progeny in response to *P. brassicae*. A) Phenotyping of F1 progeny, which exhibited a fully susceptible response (100%). Bar = 1 cm for roots panel. B) Disease phenotypes were evaluated in Col-0, F1-S, F2-R (25%), and F2-S (75%) plants using a 0–8 severity scale. A Kruskal–Wallis test was first used to assess differences among ecotypes. Pairwise comparisons were then performed using the Wilcoxon rank-sum test (Chi-square = 156.99, df = 3, P-value < 2.2e-16). Statistical analyses were conducted in R (version 4.4.2) and RStudio (version 2024.12.1+563) using the packages stats, survival, and graphics, with visualizations generated using ggplot2.

This study expands the landscape of known clubroot resistance in *A. thaliana*, identifying new ecotypes and uncovering unexpected patterns of RPB1 expression. The discovery of DraII-6 as a highly resistant ecotype, coupled with its unique early RPB1 expression profile, challenges current assumptions about the gene’s function and regulation. These findings not only provide a deeper mechanistic understanding of host–pathogen interactions but also emphasize the complexity behind durable resistance. Additionally, the finding that CR follows a recessive inheritance pattern in this ecotype opens the door to exploring other potential CR genes. Future efforts should prioritize the functional dissection of DraII-6’s resistance, including transcriptomic and epigenetic profiling, to identify additional factors that may interact or cooperate with RPB1 and play key roles in the regulation of clubroot resistance (Gravot et al., 2024). Ultimately, insights gained from this model may inform the development of more resilient Brassica crops in the face of an evolving pathogen landscape.

## ACKNOWLEDGEMENTS

We thank Dr. Abraão Almeida Santos for figure preparation and statistical analysis, and Dr. Marina Silvestre Vañó for organizing the ecotypes and providing training on Arabidopsis cultivation. Melaine Gonzalez-Garcia acknowledges the Faculty of Agricultural and Food Sciences (FSAA) at Université Laval for the *Citoyen et citoyenne du monde* – volet excellence scholarship, as well as the Fonds de recherche du Québec – Nature et technologies (FRQNT) for PhD scholarship support.

## DATA AVAILABILITY STATEMENT

All ecotypes used in this study are available from the Arabidopsis Biological Resource Center (ABRC, hZps://abrc.osu.edu/), and all accession numbers are listed in Table S1.

## SUPPORTING INFORMATION

**Figure S1**. Distribution map of the 60 ecotypes assessed in this study.

**Figure S2**. PCR amplification of *RPB1* in S, IR, and R ecotypes.

**Figure S3**. Relative expression of RPB1 in DraII-6, Est-1, and RLD-1 at 4- and 7 dpi.

**Table S1**. Ecotypes used in this study and ABRC accession numbers.

**Table S2**. Criteria used to determine the disease index based on infection severity.

**Table S3**. Primers used in this study to quantify *P. brassicae* titer, assess RPB1 presence in ecotypes, and analyze RPB1 expression.

**Table S4**. Disease index and pathogen load presented in Figure 1A, as well as the percentage of classes recorded for each CR ecotype during phenotyping (data shown in Figure 1B).

**Table S5**. Presence or absence of RPB1 in the 60 ecotypes studied here, with comparisons to other studies when data were available.

**Table S6**. Classification of CR ecotypes based on survival or death between 7 and 42dpi.

**Table S7**. RPB1 expression in Est-1, DraII-6, and RLD-1.

**Table S8**. Percentage of disease classes recorded for Col-0, the F1 progeny of a DraII-06 × Col-0 cross, and the F2 progeny obtained by open pollination of F1 plants phenotyped as resistant (25%) and susceptible (75%)

## REFERENCES

Alamery, S., Tirnaz, S., Bayer, P., Tollenaere, R., Chaloub, B., Edwards, D. et al. (2018) Genome-wide identification and comparative analysis of NBS-LRR resistance genes in Brassica napus. Crop & Pasture Science, 69, 72–93.

Alix, K., Lariagon, C., Delourme, R. & Manzanares-Dauleux, M.J. (2007) Exploiting natural genetic diversity and mutant resources of Arabidopsis thaliana to study the A. thaliana Plasmodiophora brassicae interaction. Plant Breeding, 126, 218–221.

Cao, T., Rennie, D.C., Manolii, V.P., Hwang, S.F., Falak, I. & Strelkov, S.E. (2014) Quantifying resistance to Plasmodiophora brassicae in brassica hosts. Plant Pathology, 63, 715–726.

Deora, A., Gossen, B.D. & McDonald, M.R. (2013) Cytology of infection, development and expression of resistance to Plasmodiophora brassicae in canola. Annals of Applied Biology, 163, 56–71.

Dixon, G.R. (2009) The occurrence and economic impact of Plasmodiophora brassicae and clubroot disease. Journal of Plant Growth Regulation, 28, 194–202.

Fuchs, H. & Sacristan, M.D. (1996) Identification of a gene in Arabidopsis thaliana controlling resistance to Clubroot (Plasmodiophora brassicae) and characterization of the resistance response. Molecular Plant-Microbe Interactions, 9, 91.

Glander, S., He, F., Schmitz, G., Witten, A., Telschow, A. & de Meaux, J. (2018) Assortment of Flowering Time and Immunity Alleles in Natural Arabidopsis thaliana Populations Suggests Immunity and Vegetative Lifespan Strategies Coevolve. Genome Biology and Evolution, 1, 10(9).

Gravot, A., Liégard, B., Quadrana, L., Veillet, F., Aigu, Y., Bargain, T., Bénéjam, J., Lariagon, C., Lemoine, J., Colot, V., Manzanares-Dauleux, M.J. & Jubault, M. (2024) Two adjacent NLR genes conferring quantitative resistance to clubroot disease in Arabidopsis are regulated by a stably inherited epiallelic variation. Plant Communications, 13, 5(5).

Javed, M.A., Schwelm, A., Zamani-Noor, N., Salih, R., Silvestre Vañó, M., Wu, J. et al. (2023) The clubroot pathogen Plasmodiophora brassicae: aprofile update. Molecular Plant Pathology, 24, 89–106.

Javed, M.A., Mukhopadhyay, S., Normandeau, E., Brochu, A.-S., & Pérez-López, E. (2024). Telomere-to-telomere genome assembly of the clubroot pathogen Plasmodiophora brassicae. Genome Biology and Evolution, 16(6), evae122.

Jubault, M., Lariagon, C., Simon, M., Delourme, R. & Manzanares-Dauleux, M.J. (2008) Identification of quantitative trait loci controlling partial clubroot resistance in new mapping populations of Arabidopsis thaliana. Theoretical and Applied Genetics, 117, 191–202.

Malinowski, R., Smith, J. A., Fleming, A. J., Scholes, J. D. & Rolfe, S.A. (2012) Gall formation in clubroot-infected Arabidopsis results from an increase in existing meristematic activities of the host but is not essential for the completion of the pathogen life cycle. The Plant Journal, 71, 226–238.

Malinowski, R., Truman, W. & Blicharz, S. (2019) Genius architect or clever thief – how Plasmodiophora brassicae reprograms host development to establish a pathogen-oriented physiological sink. Molecular Plant-Microbe Interactions, 32, 1259–1266.

Mukhopadhyay, S., Garvetto, A., Neuhauser, S. & Pérez-López, E. (2024) Decoding the Arsenal: Protist Effectors and Their Impact on Photosynthetic Hosts. Molecular Plant-Microbe Interactions, 37(6), 498–506.

Mukhopadhyay, S., Javed, M.A., Wu, J., & Pérez-López, E. (2025). Structure-guided secretome analysis of gall-forming microbes offers insights into effector diversity and evolution. eLife, 14, e105185.

Ochoa, J. C., Mukhopadhyay, S., Bieluszewski, T., Jedryczka, M., Malinowski, R. & Truman, W. (2023) Natural variation in Arabidopsis responses to Plasmodiophora brassicae reveals an essential role for Resistance to Plasmodiophora brasssicae 1 (RPB1). The Plant Journal, 116:1421–1440.

Peng, G., Lahlali, R., Hwang, S.-F., Pageau, D., Hynes, R.K., McDonald, M.R. et al. (2014) Crop rotation, cultivar resistance, and fungicides/biofungicides for managing clubroot (Plasmodiophora brassicae) on canola. Canadian Journal of Plant Pathology, 36, 99–112.

Salih, R., Javed, M.A., Brochu, A.S., Prakash, P., Côté, J.D. & Pérez-López, E. (2024) A Basic Guide to the Propagation and Manipulation of the Clubroot Pathogen, Plasmodiophora brassicae. Current Protocols, 4(4):e1039.

Sedaghatkish, A., Gossen, B.D., Yu, F., Torkamaneh, D. & McDonald, M.R. (2019) Whole-genome DNA similarity and population structure of Plasmodiophora brassicae strains from Canada. BMC Genomics, 20, 744.

Storfie, E., Manolii, V. P., Aigu, Y., Botero-Ramirez, A., Harding, M. W., Hwang, S. F. & Strelkov, S. E. (2025) Collection and identification of Plasmodiophora brassicae pathotypes from western Canada in 2021–2023. Canadian Journal of Plant Pathology, 1–12.

Strelkov, S.E., Hwang, S.F., Manolii, V.P., Cao, T., Fredua-Agyeman, R., Harding, M.W. et al. (2018) Virulence and pathotype classification of Plasmodiophora brassicae populations collected from clubroot resistant canola (Brassica napus) in Canada. Canadian Journal of Plant Pathology, 40, 284–298.

Vañó, M.S., Nourimand, M., MacLean, A., Pérez-López, E. (2023) Getting to the root of a club - Understanding developmental manipulation by the clubroot pathogen. Seminars in Cell Developmental Biology, 148-149, 22–32.

Wang, W., Qin, L., Zhang, W., Tang, L., Zhang, C., Dong, X. et al. (2023) WeiTsing, a pericycle-expressed ion channel, safeguards the stele to con-fer clubroot resistance. Cell, 186, 2656–2671.

Yang, Z., Jiang, Y., Gong, J., Li, Q., Dun, B., Liu, D. et al. (2022) R gene triplication confers European fodder turnip with improved clubroot resistance. Plant Biotechnology Journal, 20, 1502–1517.

