## Supplementary Figures for "Arabidopsis ecotype screening reveals novel sources of clubroot resistance and insights into resistance inheritance"

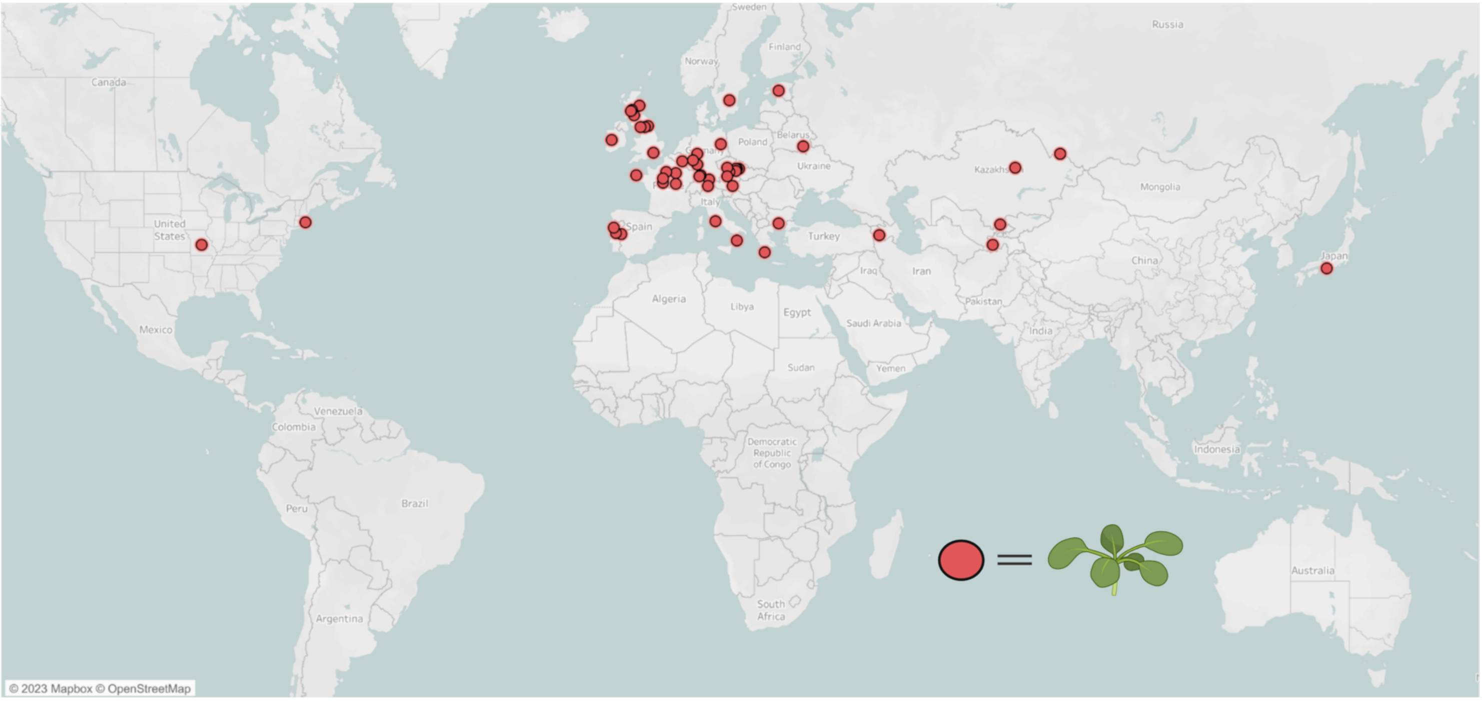


**Figure S1.** Distribution map of the 60 ecotypes assessed in this study.

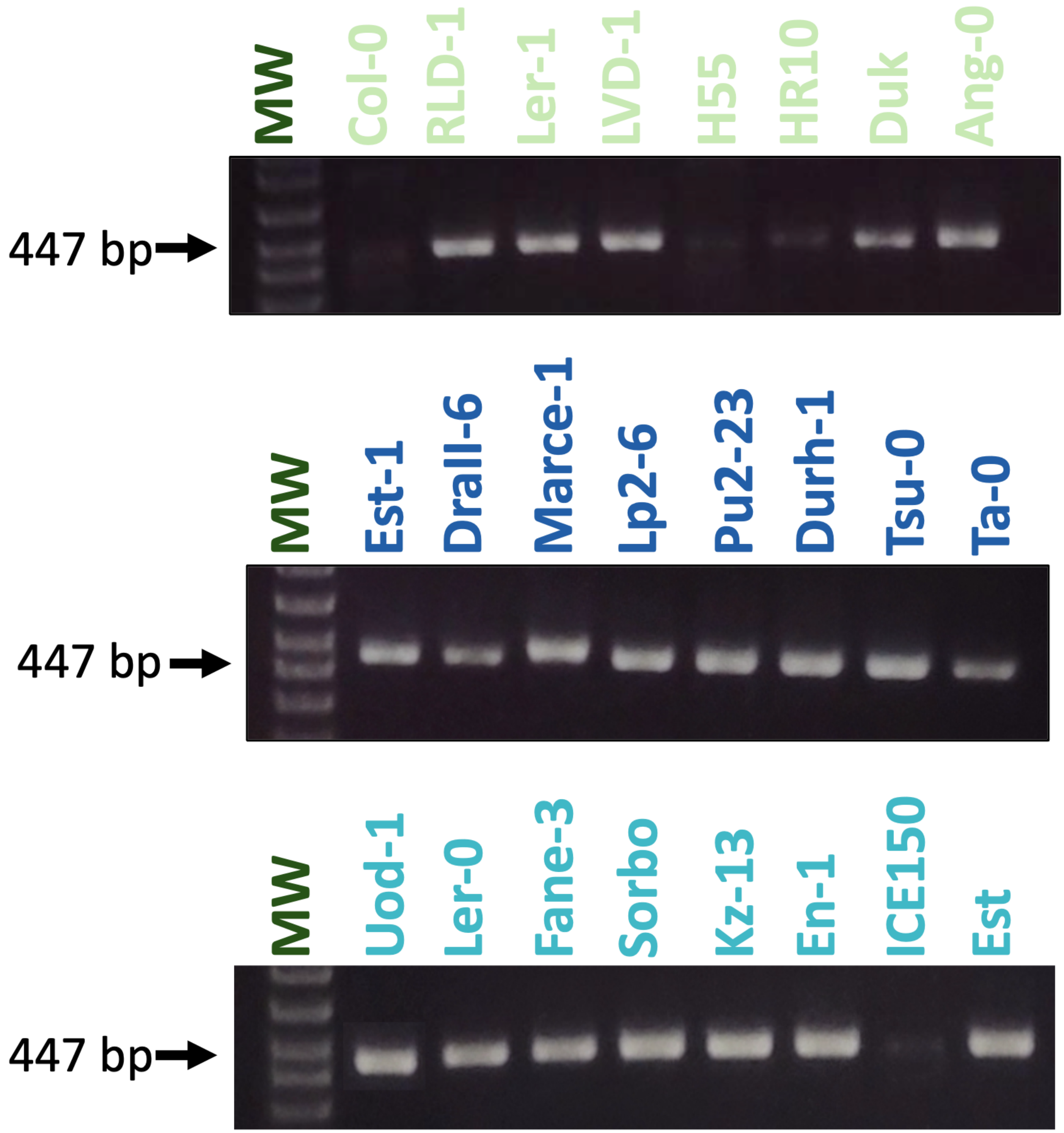


**Figure S2.** PCR amplification of *RPB1* in susceptible, intermediate-resistant, and resistant ecotypes. MW: molecular weight ladder, 1 kb Plus (Thermo Fisher Scientific, Canada).

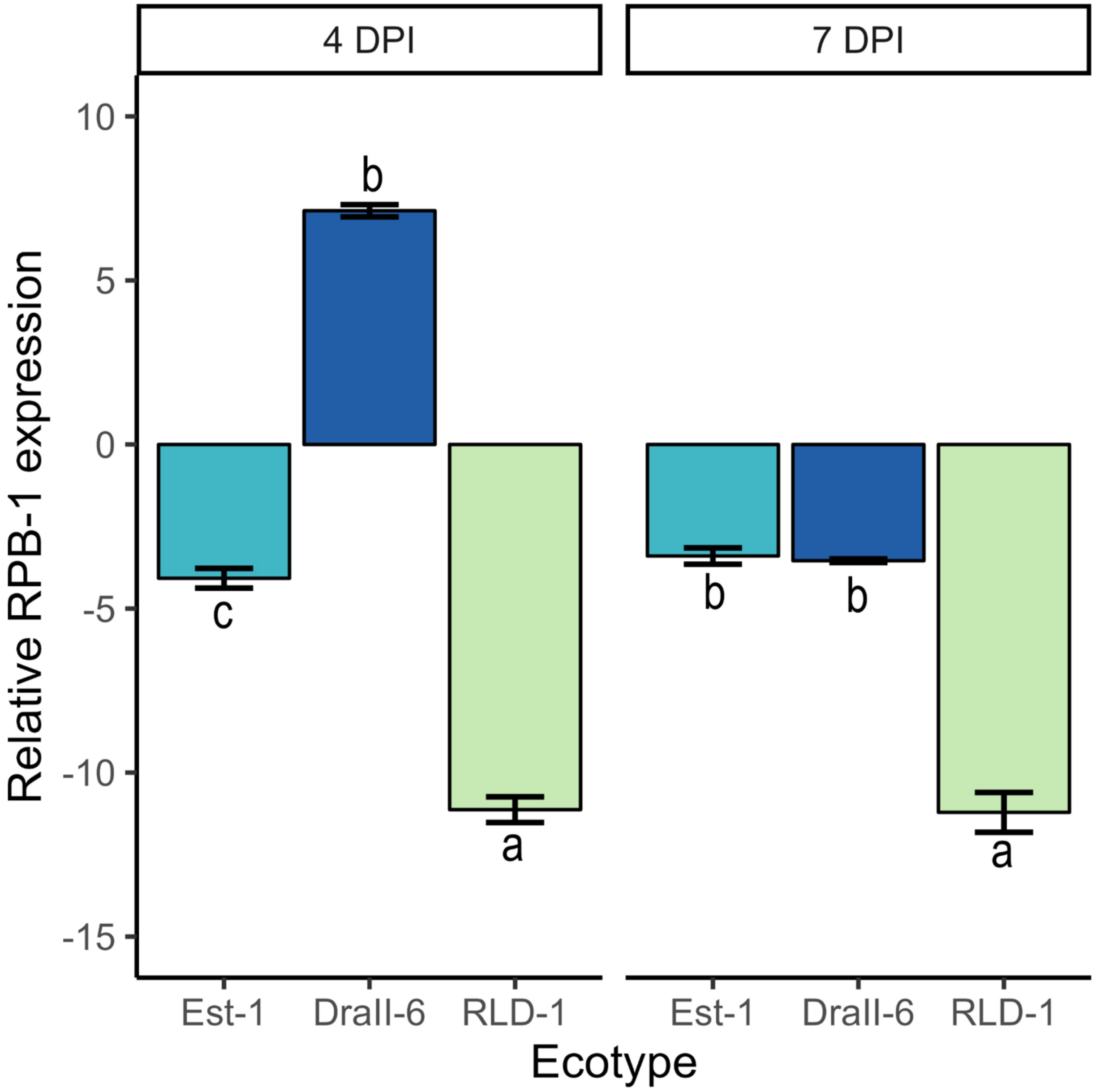


**Figure S3.** Relative expression of RPB1 in DraII-6, Est-1, and RLD-1 at 4- and 7 dpi. An analysis of variance (ANOVA) was performed separately for each time point to assess differences among ecotypes. Assumptions of residual normality and homogeneity of variances were verified. A Tukey's HSD test was then used to compare means among ecotypes at each time point. Because relative expression values were negative and no standard probability distribution exists for negative values, the analysis was conducted using their absolute positive values. At 4 dpi: ANOVA, $F = 134.95$; $df = 2, 6$; $P < 0.0001$. At 7 dpi: ANOVA, $F = 138.20$; $df = 2, 6$; $P < 0.0001$.
